# Fetal whole-heart 4D flow cine MRI using multiple non-coplanar balanced SSFP stacks

**DOI:** 10.1101/635797

**Authors:** Thomas A. Roberts, Joshua FP van Amerom, Alena Uus, David FA Lloyd, Anthony N. Price, Jacques-Donald Tournier, Laurence H. Jackson, Shaihan J Malik, Milou PM van Poppel, Kuberan Pushparajah, Mary A Rutherford, Reza Rezavi, Maria Deprez, Joseph V. Hajnal

**Affiliations:** School of Biomedical Engineering & Imaging Sciences, King’s College London, London, SE1 7EH, UK; Department of Congenital Heart Disease, Evelina Children’s Hospital, London, SE1 7EH, UK; Centre for the Developing Brain, King’s College London, London, SE1 7EH, UK

**Keywords:** cardiac MRI, fetal imaging, SSFP, congenital heart disease, flow, velocity, 4D reconstruction

## Abstract

**Purpose:** To develop an MRI framework for reconstruction of 4D velocity vector blood flow volumes for visualisation and quantification of circulation in the fetal heart and major vessels.

**Methods:** A novel method of velocity-encoding using multiple non-coplanar stacks of bSSFP phase images was combined with a previous framework for reconstruction of motion-corrected 4D magnitude cine volumes to generate spatiotemporally-paired 4D flow cine volumes of the fetal circulatory system. The multiple stack velocity-encoding scheme was validated in a simulated flow phantom and compared with a gold-standard method for velocity-encoding in a physical flow phantom. The 4D flow cine framework was evaluated in seven fetal subjects. Reconstructed 4D flow volumes were evaluated by an expert fetal cardiologist and preliminary flow measurements were taken in various major vessels of the heart.

**Results:** Phantom experiments showed that the multiple non-coplanar stack velocity-encoding scheme was accurate. The 4D flow cine reconstruction framework was robust in fetal subjects and generated multi-dimensional velocity vector maps of blood flow through the cardiac cycle. Directionality of blood flow was consistent with expected fetal circulatory hemodynamics. Relative blood flow rates in the major vessels were in line with previous observations, although absolute values were underestimated by a factor of approximately two due to limitations of spatial and temporal resolution.

**Conclusion:** 4D flow cine volumes can be reconstructed from multiple non-coplanar stacks of slices. The proposed framework was used to visualise and quantify flow through the whole fetal heart and great vessels, but is applicable to any imaging scenario where motion is a major challenge.

## 1 INTRODUCTION

Blood flow imaging of the fetal heart and great vessels is extremely challenging because the heart beats very rapidly, the vessels are small and the fetus is prone to spontaneous movement as well as displacements caused by maternal respiration. Pulsed wave Doppler echocardiography^1–3^ is readily available in clinics for measuring blood flow rates, and is very low cost compared to using MRI, but in practice is rarely used in fetal or paediatric cardiology. It relies on line-of-sight velocity assessment combined with assumptions about the shape and the flow profile of the targeted blood vessel to convert measurements to flow rates, and can only provide data from a single site at a time. Further still, the accuracy, precision and reliability of measurements also depend on fetal lie, maternal habitus and the level of expertise of the operating sonographer^4^.

Phase contrast (PC) MRI methods are routinely used for time-resolved quantitative blood flow measurements of the cardiovascular system in adult hearts^5–7^, and to a lesser extent in paediatric populations^8–10^ and in neonates^11,12^. In the last decade, some developments have been made towards using PC-MRI for imaging flow in the fetal cardiovascular system^13–15^, however, these approaches have been limited to single-slice acquisition, which can be heavily compromised in the event of fetal motion. Minor fetal motion, such as small displacements compared to the target vessel, can compromise the accuracy of velocity measurements, and at worst, major fetal motion can shifting the vessel out of the imaging plane entirely. This can lead to a scenario where the operator must repeatedly perform new, manually specified pilot scans, then re-acquire the PC-MRI scan until the desired images are obtained. This can be time-consuming and challenging given limited available examination time and the size of the anatomy. Moreover, even when successfully achieved, two-dimensional imaging is intrinsically limited given the wide range of anatomical arrangements and size of the heart and great vessels in congenital heart disease. Three-dimensional imaging permits visualisation of the intricate fetal vasculature and the complex connections and shunts within the developing heart. 4D flow imaging has been very recently demonstrated in the sheep fetus^16^, albeit with the use of anaesthetic to immobilise the mother and fetus, but until now only single-slice methods have been carried out *in utero* in the human fetus.

Recently, we have developed an MRI framework for motion-tolerant 3D imaging of the fetal cardiovascular system based upon the imaging principle of acquiring multi-planar stacks of slices for volumetric reconstruction with slice-to-volume registration (SVR), originally developed for fetal brain imaging^17–20^ and have demonstrated its clinical utility^21^. This was then extended to time-resolved 4D cine imaging of the fetal heart^22,23^. These approaches dispense with the need for precise single-slice planning; instead the operator is simply required to cover the fetal heart with a sufficient number of non-coplanar stacks. The final reconstructed 4D cine volumes allow for interpretation of the complicated fetal cardiac vessel structures and intra-chamber connections, in any 2D plane.

In this paper, the volumetric 4D cine framework was extended to generate *in utero*, motion-corrected, time and 3D spatially-resolved full vectoral velocity distributions of blood flow in the whole fetal heart and great vessels. In the previous framework, 4D cine volumes were generated using a k-t SENSE^24,25^ accelerated balanced steady-state free precession (bSSFP) multi-planar real-time acquisition combined with retrospective image-domain techniques for motion correction and cardiac gating. Here, velocity-encoding is achieved by exploiting the inherent flow sensitivity of bSSFP images. This has been demonstrated and exploited previously by several authors who have devised methods to achieve classic PC-MRI velocity mapping by modifying the imaging gradients^26,27^. As bSSFP sequences possess an intrinsic velocity sensitivity that has a fixed (although oblique) direction relative to the imaging plane, the diversely-oriented planes which are a feature of SVR reconstructions provide multiple non-colinear sensitisations that could be used to recover full velocity information. In this paper, we explore this concept and present an initial framework for recovering fully motion corrected 4D velocity fields in the fetal heart. The methods deployed build on the previous fetal cine MRI framework^23^ and a scattered data SVR approach adapted from fetal diffusion MRI^28^. The resulting velocity reconstruction framework was validated in both a simulated flow phantom and a physical flow phantom, and then the full 4D flow cine framework was evaluated in a cohort of seven human fetal subjects.

## 2 THEORY

When measuring a single-component of flow with a conventional phase contrast MRI sequence, such as a spoiled gradient echo (SPGR), two measurements of phase are acquired with identical sequence parameters except for different velocity-encoding gradients. Subtraction of the two phase images removes background phase offsets and phase associated with stationary tissue, allowing for calculation of the projection of any arbitrary velocity vector on to a specified spatial direction. The magnitude and direction of velocity-sensitivity is determined by the first moment (***M***) of the combined imaging and velocity-encoding gradients. For conventional three-dimensional phase contrast flow imaging, the imaging geometry is held fixed and the velocity-encoding gradients, which are usually simple bipolar trapezoids with no zeroth order moment, are switched to different axes to create a set of orthogonal velocity-sensitive imaging volumes, as shown in Figure 1a and Figure 1b.

**Figure 1:**
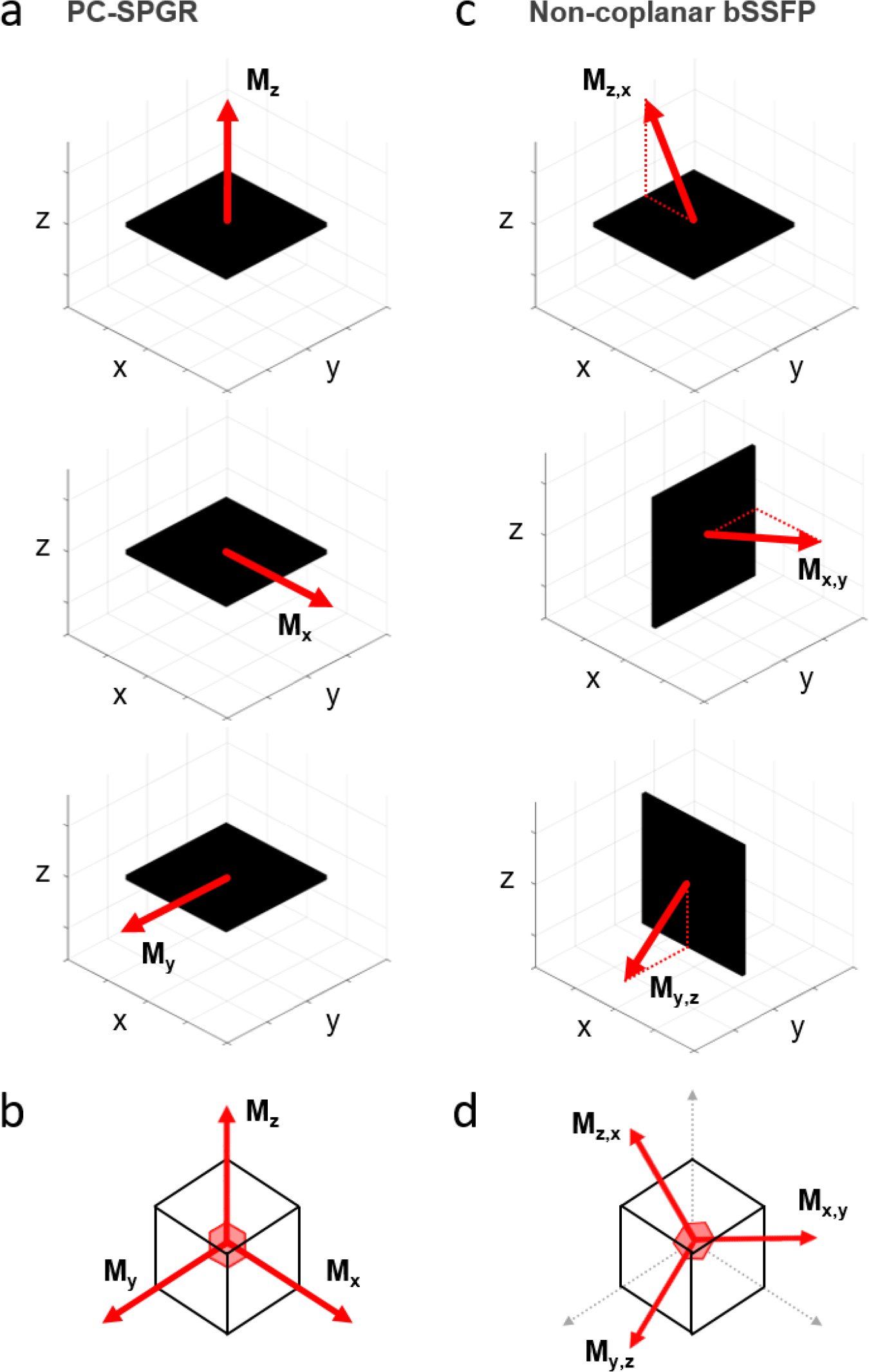
Velocity encoding using a standard spoiled gradient-echo (SPGR) paradigm versus a non-coplanar multi-stack approach compatible with slice-to-volume registration (SVR). (a) In standard velocity-encoding using dedicated sensitising gradients, the imaging volume (black) is fixed and the dedicated velocity gradient first moments (M) are played-out using different gradients (M_x_, M_y_, M_z_). Note that there are typically first order moments associated with the native imaging gradients, which add a constant additional encoding, but this is not shown as it is usually subtracted off. (b) For each imaged voxel, the three velocity-encoding directions are orthogonal and conventionally aligned with the imaging axes. (c) Volumetric reconstruction using SVR requires the acquisition of non-coplanar stacks. In this case, the directionality of the first moment associated with the intrinsic imaging gradients is maintained relative to the stack and rotation of the imaging volume provides 3D velocity-encoding. (d) With PC-bSSFP, the gradient first moment is oblique to the imaging plane, but combining three non-coplanar stacks provides data at the location of each voxel with three non-colinear velocity sensitisations, now oblique to the imaging axes.

Adding dedicated bipolar velocity encoding gradients into bSSFP sequences is challenging as these increase the repetition time (TR). However, Markl et al.^9^ devised a method for measuring through-plane flow using a bSSFP sequence by simply inverting the slice-select gradient between two consecutive acquisitions. As the readout gradients are identical between the two acquisitions, subtraction of the resulting phase images eliminates any contributions caused by all other imaging gradients. The resultant velocity-encoding is associated only with the slice-select gradient axis. Nielsen et al.^27^ proposed a time-efficient method of 3D velocity mapping using bSSFP by minimally augmenting the imaging gradients to create an additional controllable velocity sensitivity. Acquisition of three sequences with unique gradient first moments allowed for estimation of 3D flow vectors.

For fetal imaging with SVR, acquisition of multi-planar stacks in different orientations is a prerequisite for volumetric reconstruction. SVR with conventional three-dimensional velocity-encoding would be prohibitively time-inefficient because a minimum of nine acquisitions are required to sample three spatial dimensions and three velocity dimensions. Instead, we propose achieving multi-dimensional velocity-encoding by rotating stacks with identical fixed gradient first moments, as shown in Figure 1c and Figure 1d. With this scheme, a minimum of three non-coplanar stacks with (ideally orthogonal) non-colinear gradient first moments are required to sample three spatial dimensions and three velocity dimensions.

In our previous framework for fetal whole-heart 4D cine imaging, a bSSFP sequence was used for image acquisition. All three of the gradients that form bSSFP sequences with Cartesian sampling generally contribute non-zero first moments, but only the readout and slice-select gradients produce constant effects throughout the acquisition. The net effect is an oblique gradient first moment. This is directed in a plane containing the slice select and readout directions, since the effect of the phase encoding gradient, which is not zero, averages to zero. If acquisitions with three orthogonal bSSFP planes are employed, the three associated gradient moments are also orthogonal to each other (Figure 1d), but they are rotated relative to a standard 3D velocity encoding scheme (Figure 1b).

In this work, we take advantage of the diversity of imaging planes required for volumetric reconstruction of fetal hearts to sample multiple velocity-encoding directions and then to recover a full vector representation of blood flow. In theory, as in the studies by Nielsen et al.^27^, a minimum of three stacks with non-colinear first moments are required for 3D velocity-encoding, but to ensure stable inversion of phase data into velocity vectors given fetal motion, the present experiments were performed with a minimum of five bSSFP stacks.

## 3 METHODS

The proposed framework for whole-heart 4D flow cine reconstruction is shown in Figure 2. In brief, the reconstruction consists of two streams: first, motion corrected 4D magnitude cine reconstruction is performed adhering to the pipeline outlined previously described by van Amerom et al.^23^. The data consists of multiple stacks of real-time bSSFP imaging planes, with multiple dynamics (frames) obtained for each slice. The fetal heart rate is estimated from the frames of each acquired slice and used to assign cardiac phases to each successive frame. The cardiac phases of different slice locations are synchronised and then rigid body transformations are determined for each individual frame to provide a fully motion corrected dataset for reconstruction.

**Figure 2:**
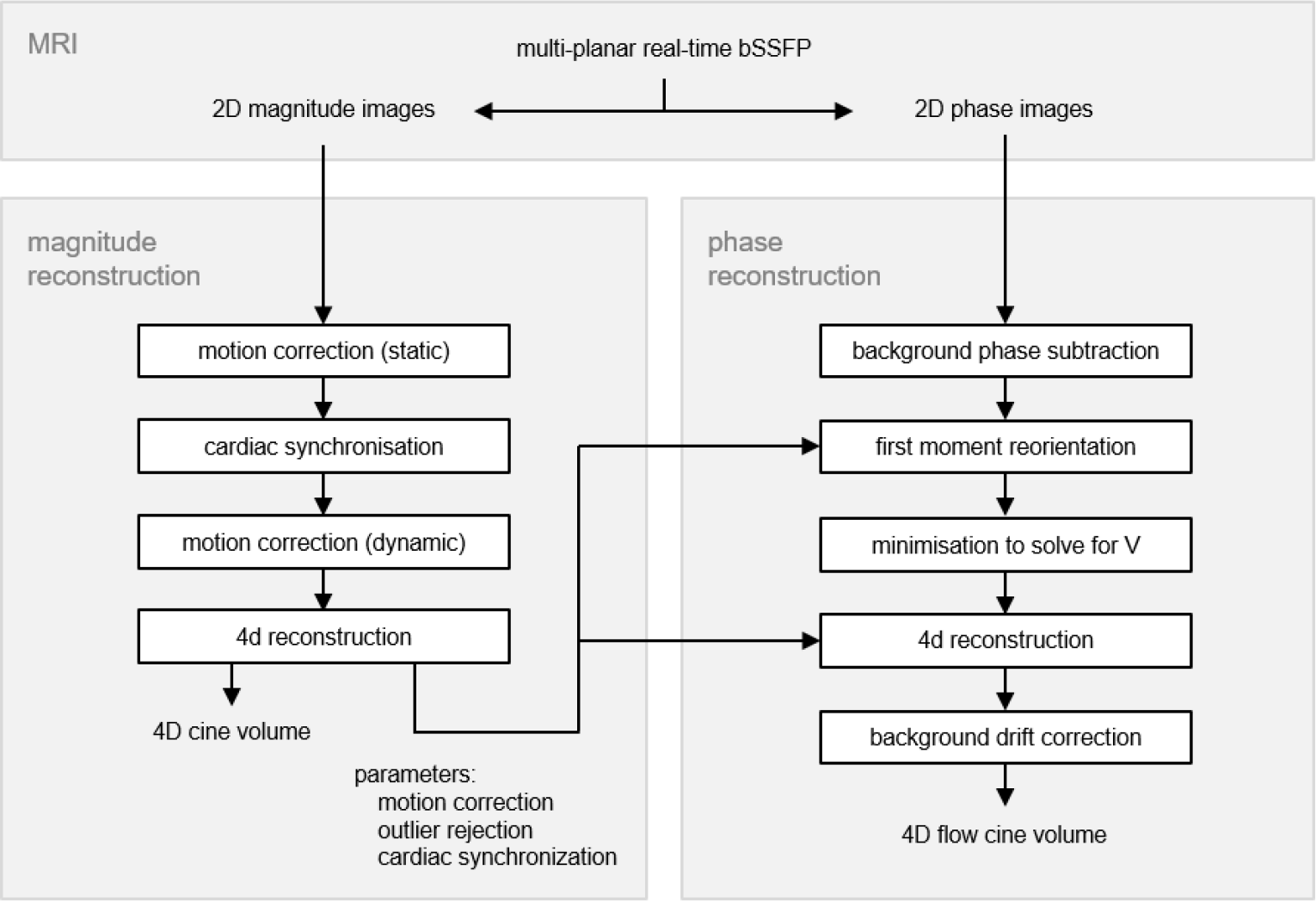
Framework for 4D vector flow cine volume reconstruction. Top panel: multi-planar real-time accelerated bSSFP data is acquired and reconstructed into 2D magnitude and phase images using k-t SENSE. Left panel: volumetric 4D magnitude cine reconstruction of the fetal heart is performed, as detailed previously^23^. Right panel: volumetric 4D vector flow cine reconstruction of the fetal heart is performed, using motion correction, cardiac synchronization and frame outlier rejection parameters from the magnitude reconstruction pathway. Background phase is estimated and subtracting by polynomial approximation. First moment vectors for the imaging gradients, as specified for the native acquisitions, are rotated as a result of detected motion. Velocity vector fields are recovered by inversion of the phase image information using a conjugate gradient method. Volumetric and temporal 4D reconstruction is performed before a final background velocity drift correction is applied.

Parameters generated from this pipeline, including stack and frame spatial transformations, motion correction parameters and cardiac synchronisation parameters are then passed to the phase data reconstruction stream. Background phase subtraction, gradient moment reorientation and calculation of velocity volumes are performed, before a final velocity drift correction is applied. The complete framework results in coupled 4D magnitude cine and 4D flow cine volumes which are spatially and temporally equivalent. For consistency, the mathematical notation in this manuscript follows the same logic and conventions used in the 4D magnitude cine paper^23^.

### 3.1 Multi-Planar Dynamic MRI

Multi-planar, dynamic MR images were acquired in stacks of parallel slices with 96 dynamics per slice position and reconstructed using k-t SENSE^24^ (acceleration factor 8) with regularisation, in an identical manner to that described previously^23^ (Figure 3a). Stacks were acquired in multiple orientations to ensure full coverage of the heart and the great vessels. The acquired data form a set of *N*_*k*_ dynamic MR image frames of complex type, 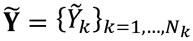, where each frame, k, has an acquisition time *t*_*k*_ and consists of elements 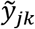 at 2D spatial coordinates indexed by *j*. The complex data is reconstructed into magnitude image frames, 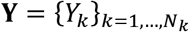, and phase image frames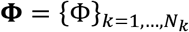.

**Figure 3:**
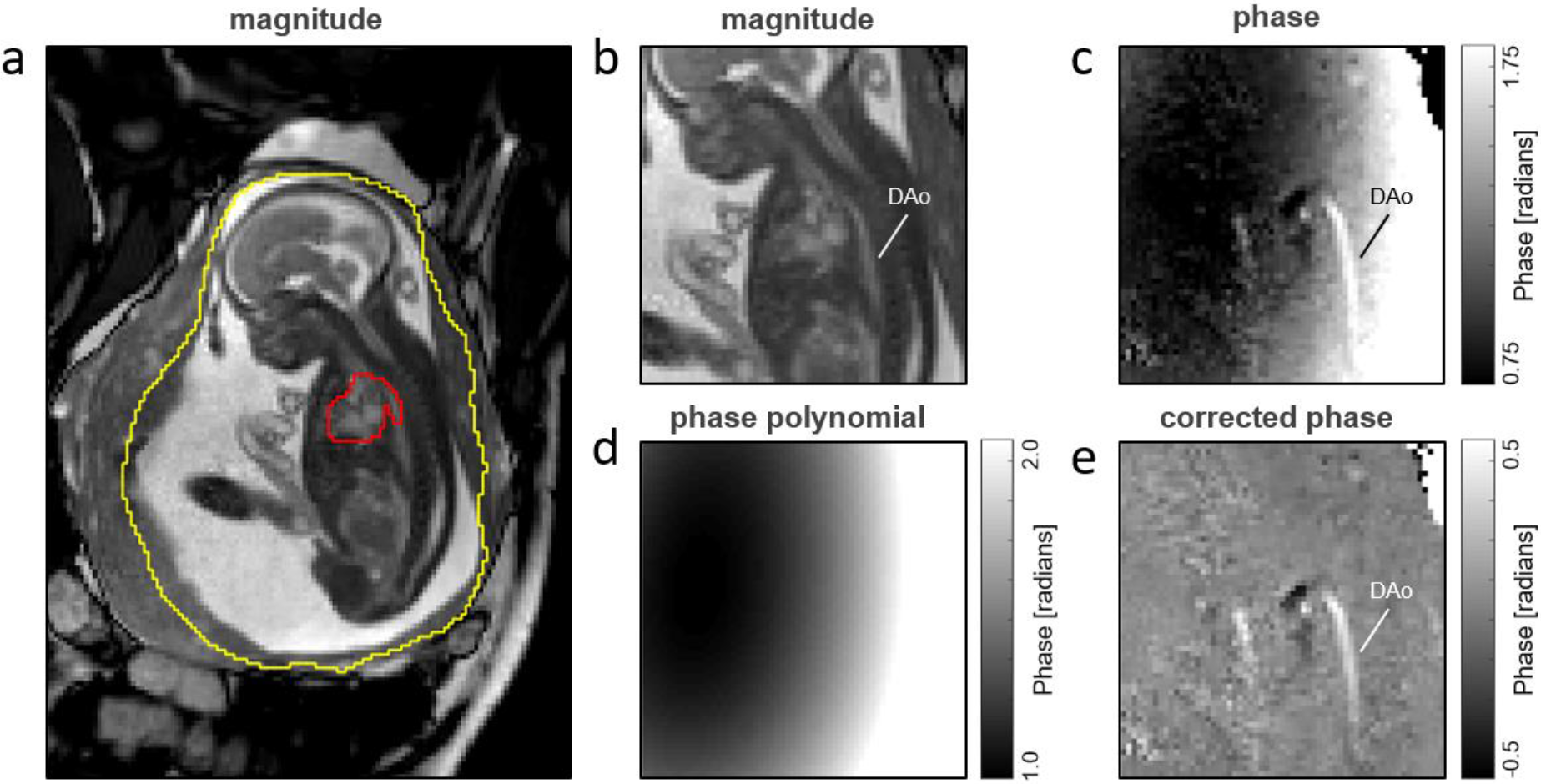
Background phase correction of bSSFP images shown in a healthy 24^+2^ week fetus (ID03). (a) Magnitude image showing two (multi-planar) regions of interest drawn around the mother’s uterus (yellow) and the fetal heart, including great vessels (red). (b) Zoomed-in magnitude image showing the descending aorta (DAo). (c) Uncorrected phase image demonstrating the background phase variation. (d) Third-order polynomial fit calculated from the uncorrected phase image using the uterus region of interest but excluding the fetal heart and great vessels. (e) Corrected phase image after subtraction of the polynomial phase estimate leaving localised phase changes that are then used to calculate the velocity vector field.

### 3.2 4D Magnitude CINE Reconstruction

The magnitude image frames, **Y**, were used for reconstruction of a 4D magnitude cine volume 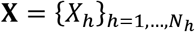, where X_*h*_ has elements *X*_*ih*_ for spatial index *i* and temporal index *h* corresponding to cardiac phase, 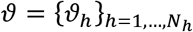, as described in our previous framework^23^. The magnitude pipeline also includes an outlier detection process that operates at the voxel and frame level and is used to reduce the weight of discordant signals that are likely to be causes by image artefacts. The spatial locations of voxels in the magnitude images were identical to those in the phase image frames, ***Φ***, as both image types were generated from the same bSSFP acquisition. Therefore, reconstruction parameters and weights from the 4D magnitude cine stream could be passed to the 4D flow cine reconstruction stream. These included:

1. Rigid body transformation matrices, **A**, which align dynamic image frames with the output cine volume.
2. Image frame outlier rejection weights, **P**_**k**_, which exclude or include the specified frames from further processing.
3. Cardiac cycle parameters 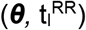, which map acquired image frames with phases of the cardiac cycle.

### 3.3 Background Phase Correction

The phase of each bSSFP image is offset by variation in the main magnetic field, as visible in Figure 3c. This background phase must be eliminated to accurately estimate the motion-induced phase changes within the heart. After some initial experimentation, the background phase offset was estimated in each stack by fitting a third-order polynomial^27^ (Figure 3d) to the phase images using a manually drawn multi-planar region of interest (Figure 3a) which included the relatively static contents of the uterus (yellow), whilst excluding the rapidly-moving heart (red). Subtraction of the polynomial resulted in corrected phase images related to the underlying velocity, with minimal remaining background phase variation (Figure 3e).

### 3.4 Gradient Moment Correction

Every slice within a single stack of phase data is acquired with the same configuration of gradient first moments, which corresponds to a fixed velocity-encoding direction relative to the slice. If fetal and/or maternal motion causes the slice to be rotated in anatomical space, then the gradient first moment is subject to the same rotation. However, the rotated slice still provides a valid projection of the velocity given the change in gradient first moment. In the proposed framework, the gradient first moments of each frame are recalculated by applying the same slice transformation matrices (**A**) that were applied to the equivalent slices in the 4D magnitude cine reconstruction.

### 3.5 4D Flow CINE Reconstruction

For a single bSSFP stack acquired with Cartesian sampling, the phase in a voxel, *φ*_*jk*_, is given by the dot product between the gradient first moment and local velocity (***v***_***jk***_) in the voxel:

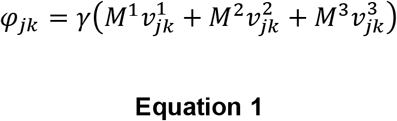

where γ is the gyromagnetic ratio, and *M*^*q*^ and *v*^*q*^_*jk*_ are the gradient first moment and velocity-components, respectively, indexed by *q* for their Cartesian components. In conventional velocity-encoding, *M*^1…3^ would typically denote the readout, phase encode and slice-select directions, but more generally *M* represents any orthonormal coordinate system. Note, that in a bSSFP sequence, the gradient moment, *M*^*q*^, associated with the phase encode direction is non-zero for each individual phase encoded data line, but for the data as a whole there is no net moment.

Fetal and maternal motion causes rotation of the gradient first moments, as described in section 3.4. Using frame-wise transformations *A*_*k*_ from the 4D cine magnitude reconstruction, frame-wise corrected gradient first moments are calculated as 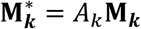 where **M_k_** is a 3-by-1 vector associated with frame, *k*, given by **M**_**k**_ = {*M*^*q*^}_*q*=1,…,3_, and **M**_**k**_* follows the same logic. The complete set of corrected gradient first moment vectors is therefore defined as matrix 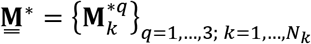 which describes the relationship between the concatenated list of three-dimensional velocity vectors at spatial location *j* and frame *k*, 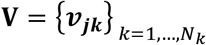, and the vectorised list of phases, ***Φ***, of all voxels according to:

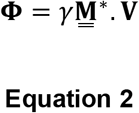

To calculate **V**, the current estimates of ***v***_***jk***_ are used to simulate the phases, 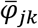, which can be subtracted from the acquired phase at the same location, *φ*_*jk*_, to obtain an error term:

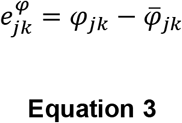

which can be minimised with by sum of squares difference using an iteratively optimised conjugate gradient descent method:

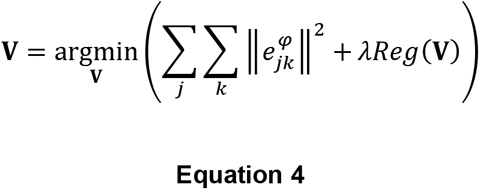

which includes a spatial edge-preserving regularisation term *Reg*(**V**) to stabilise the reconstruction and a regularisation controlling parameter, λ.

### 3.6 Velocity Drift Correction

Following reconstruction, it was observed that regions of static tissue in the 3D magnitude volumes displayed a velocity offset in the 4D cine velocity-component volumes. To correct for this drift, a third-order polynomial was fitted to the temporal median of each of the velocity-component volumes, **V**^**q**^, using a threshold mask to exclude the blood pool. Subtraction of the polynomial from each frame of the velocity-component volumes resulted in drift-corrected velocity-component images.

### 3.7 Simulated Flow Phantom Study

A simulated flow phantom was created in MATLAB for the purposes of testing and validating the reconstruction of multi-planar stacks of phase data into velocity volumes. The simulated flow phantom consisted of six cylindrical pipes arranged in pairs with antiparallel flow, orientated in three orthogonal directions. Gaussian flow profiles, which were truncated at the boundaries of the cylinders, were used as a simple approximate of laminar flow in a pipe. The pairs of pipes had different peak velocities (±25, ±80 and ±100 cm/s) chosen to simulate a range of different, physiologically-relevant flow rates. Multi-planar stacks of phase images in any orientation were simulated according to Equation 1 by using gradient first moment values taken from the scanner bSSFP acquisition. Gaussian random noise was added to the phase images so that they had a signal-to-noise ratio equivalent to the bSSFP data acquired in the physical flow phantom experiment (in section 3.8).

For the results presented, five stacks of images with 1 mm isotropic voxels were generated in different orientations: three stacks were chosen such that each was orthogonal to one set of pipes, and two stacks were specified in oblique orientations through the simulated phantom. Velocity volumes were then reconstructed using the proposed framework and the results compared with the ground truth simulated flow phantom. The simulated flow phantom MATLAB code will be made available for download upon publication of the manuscript.

### 3.8 Physical Flow Phantom Study

A simple physical constant flow phantom consisting of plastic-tubing connected to a water pump was scanned to demonstrate the proposed velocity reconstruction method. To minimise bSSFP-related artefacts, the plastic tubing was submerged in a water-filled, spherical glass flask. Two types of tubing with different internal diameters (to provide two different flow rates) were formed into a continuous loop, which returned water to the pump.

Five bSSFP stacks were acquired in the physical flow phantom: three in orthogonal orientations aligned with the scanner axes and two oblique stacks. Images were acquired with 1.5mm in-plane resolution and 3mm through-plane resolution with a 1.5mm slice overlap. With these parameters the velocity that produces a π phase shift, VENC_bssfp_, was 79 cm/s. The physical flow phantom was also imaged using a standard multi-planar PC-SPGR acquisition for comparison and validation of the proposed method. Three coplanar stacks with orthogonal velocity-encoding directions were acquired using the PC-SPGR acquisition, with voxel resolution identical to the bSSFP stacks. The PC-SPGR stacks were acquired with VENC_spgr_ = 50 cm/s. PC-bSSFP velocity volumes were reconstructed using the proposed method. PC-SPGR velocity volumes were reconstructed by standard vector addition of phase data and scaling by VENC_spgr_.

### 3.9 Fetal Study

A subset of 7 singleton pregnancies were selected from the previously published cohort^23^. The only selection criteria was the availability of 5 non-coplanar bSSFP stacks for reconstruction. This was to ensure extensive sampling of all possible directions of blood flow in the heart and to allow for redundancy needed for stable inversion, even in the event of fetal and/or maternal motion which introduces unpredictable rotations to the actual velocity encoding directions. Full details of the fetal subjects, who ranged from 24 to 33 weeks gestational age (GA), are given in Table 1. All imaging was performed with approval of the local research ethics committee and all participants gave written informed consent prior to enrolment.

**Table 1:** Fetal study subjects. Columns denote: fetal case number (ID); fetal gestational age in weeks^+days^ (GA); number of multi-planar dynamic MR stacks used for reconstruction of 4D flow cine volumes (No. Stacks).

| **ID** | **GA** | **Clinical status** | **No. Stacks** |
| --- | --- | --- | --- |
| **01** | 32<sup>+1</sup> | dilated aortic root | 5 |
| **02** | 30<sup>+6</sup> | volunteer (normal heart) | 5 |
| **03** | 24<sup>+2</sup> | volunteer (normal heart) | 5 |
| **04** | 29<sup>+6</sup> | right aortic arch | 5 |
| **05** | 31<sup>+0</sup> | ventricular diverticulum | 6 |
| **06** | 32<sup>+3</sup> | right aortic arch | 6 |
| **09** | 28<sup>+0</sup> | volunteer (normal heart) | 5 |

Sequence parameters had been optimised as described in van Amerom et al^21^ to balance good signal with high spatio-temporal resolution, while ensuring full coverage of fetal and maternal anatomy within the field of view and adhering to safety constraints. The safety constraints were whole body specific absorption rate (SAR) < 2.0 W/kg^29^, low peripheral nerve stimulation (low PNS mode in vendor software) and sound pressure level (SPL) < 85dB(A), accounting for >30dB attenuation in utero^30^. All data was acquired on a 1.5T Ingenia MRI scanner (Philips, Netherlands) using an anterior torso coil array in combination with a posterior spine coil array to measure signal in 28 receiver channels

The bSSFP sequence was run with regular Cartesian k-t undersampling^25^ with: TR/TE 3.8/1.9ms, flip angle 60°, FOV 400×304mm, voxel size 2.0×2.0×6.0mm, 8x acceleration, 72ms temporal resolution, 96 images per slice, slice overlap 2-3mm, VENC_bssfp_, = 79 cm/s. Coil calibration data were acquired prior to bSSFP acquisition and k-t training data were acquired following the under-sampled acquisition. Acquisition time of a single stack was typically 155s.

k-t SENSE reconstructon of bSSFP data was performed in MATLAB (Mathworks, USA), with additional functionality from ReconFrame 3.0.535 (GyroTools, Switzerland). The 4D flow cine reconstruction framework was implemented using a combination of MATLAB and both the Image Registration Toolkit (IRTK, BioMedIA, UK) and the Medical Image Registration Toolkit (MIRTK, BioMedIA, UK), extending work by Kuklisova et al.^17^, van Amerom et al.^22,23^ and Deprez et al.^28^. In keeping with the 4D magnitude cine framework, 4D flow cine volumes were reconstructed with an isotropic spatial resolution of 1.25mm and *N*_*h*_ = 25 cardiac phases. The code underlying the proposed framework will be made available for download upon publication of the manuscript.

### 3.10 Evaluation of 4D Flow CINE Volumes

Whole-heart 4D flow cine volumes for each subject were assessed by an expert MRI fetal cardiologist (DL) using MRtrix3^31^ to allow visualisation in any orientation with optional overlay of velocity vectors on the anatomical cine images. For the purpose of calculating temporal blood flow curves, single cross-sectional 2D regions of interest were manually drawn perpendicular to selected blood vessels using the Medical Imaging Interaction Toolkit (MITK) Workbench software (German Cancer Research Center, Germany) which allowed free-rotation of the 4D magnitude cine volume reconstruction. The following vessels were sampled: ascending aorta (AAo), descending aorta (DAo), pulmonary artery (PA), superior vena cava (SVC) and ductus arteriosus (DA). Voxel-wise blood flow was calculated as the product of velocity vector magnitude 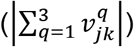 and voxel area (1.25 mm^2^). Vessel blood flow was calculated as the sum of all voxels within the region of interest. Finally, these measurements were normalised to gestational weight based on previously published fetal growth curves^32^.

## 4 RESULTS

Results from the simulated flow phantom are shown in Figure 4c. The reconstructed velocity volume closely matched the original synthetic flow phantom. The difference image between the reconstructed velocity volume and the synthetic flow phantom (Figure 4d) in regions corresponding to the pipes was within the level of the noise, except for regions which were not intersected by all five phase stacks (black arrow on difference image). This vulnerability to insufficient sampling motivated the use of 5 stacks for the in vivo data.

**Figure 4:**
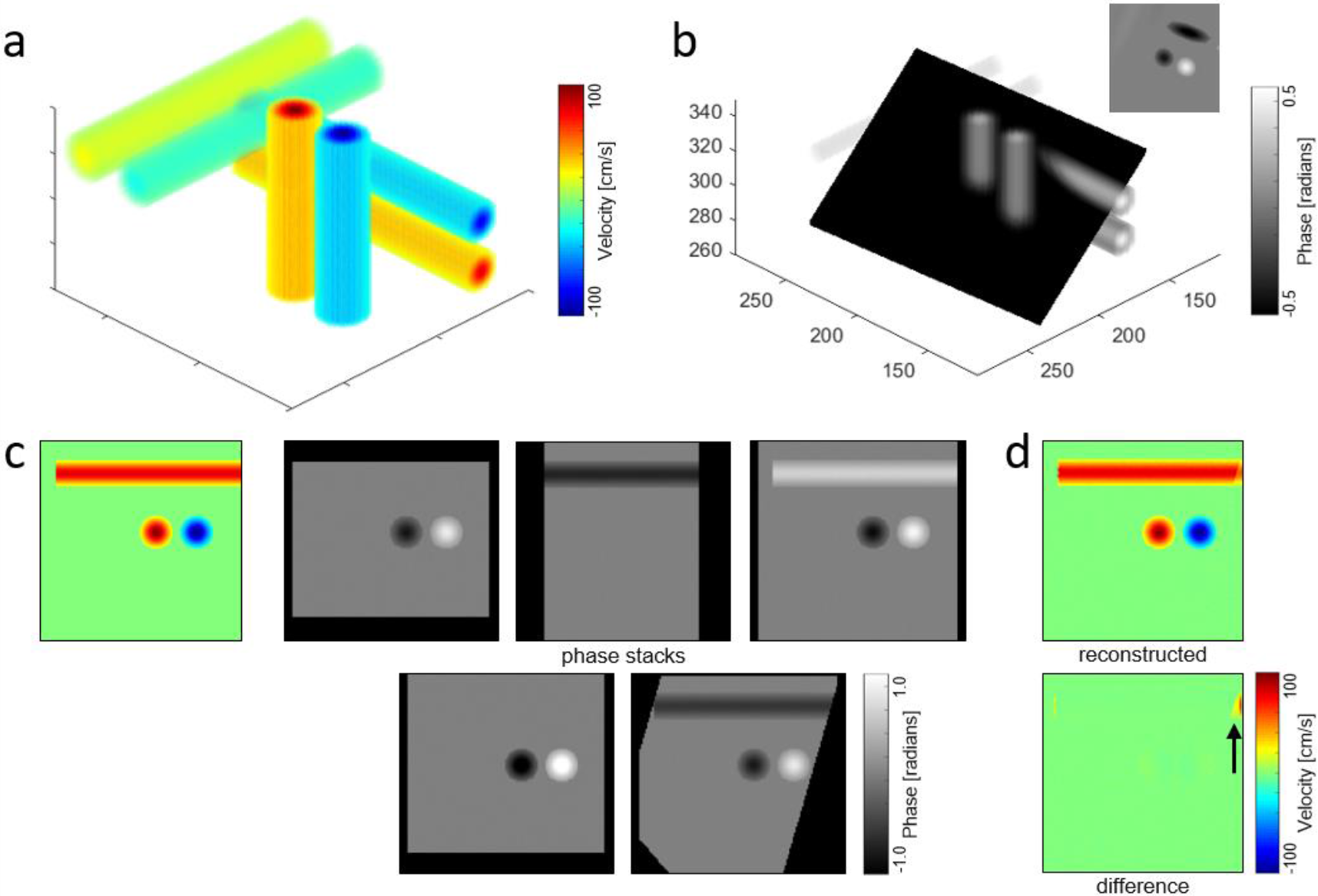
Simulated flow phantom experiment. (a) The phantom consisted of six orthogonal pipes with variable flow velocities (±25, ±80, ±100 cm/s). (b) Example of a possible oblique slice projection through the phantom and, inset, the corresponding in-plane phase image. (c) Ground truth flow velocities (colour) and phase images (greyscale) from five non-coplanar stacks, all at the same slice location. (d) 3D velocity reconstruction at an equivalent location to and the corresponding subtraction image to show reconstruction errors. Black arrow shows region not covered by all phase stacks.

Cross-sectional magnitude images of the physical flow phantom are shown in Figure 5a and Figure 5b. Figure 5c shows reconstructed velocity components from within a region of the flow phantom which was covered by all five bSSFP stacks. The directionality and the speed of the flowing water was consistent between the bSSFP and the PC-SPGR velocity volume reconstructions. Velocity profiles through pipes of different internal diameters demonstrated strong agreement between the PC-SPGR and PC-bSSFP methods, as shown in Figure 5d. The peak velocity magnitude of the profile across pipe 1 (P1) was V_SPGR_ = 37 cm/s and V_bSSFP_ = 34 cm/s. In pipe 2 (P2) the peak velocity magnitude values were V_SPGR_ = 13 cm/s and V_bSSFP_ = 12 cm/s.

**Figure 5:**
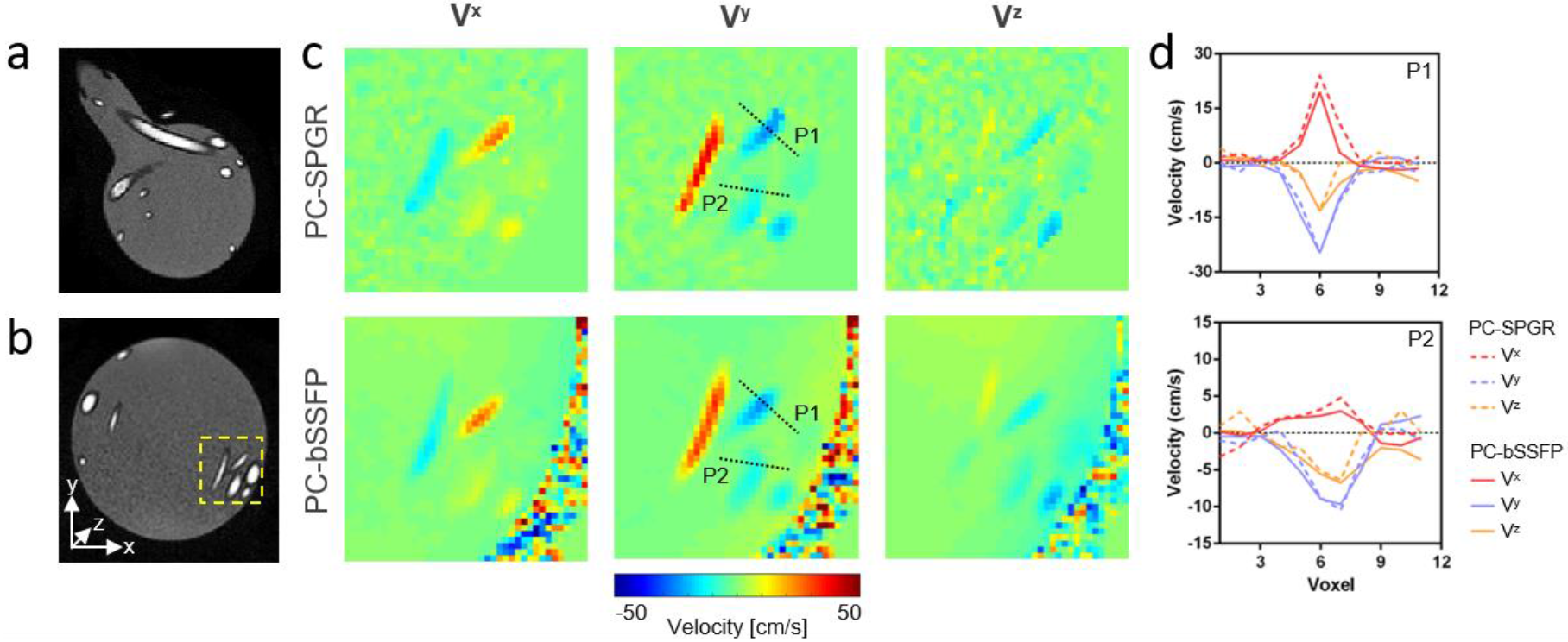
Physical flow phantom experiment. (a), (b) Magnitude images of the physical flow phantom, which consisted of a spherical water filled glass flask containing tubes connected to a flow pump. (c) Measured velocity components from the region of interest (dashed yellow) shown in (b). Top row shows velocity components from a PC-SPGR acquisition reconstructed using the standard scanner vendor processing. Bottom row shows velocity components from a bSSFP acquisition reconstructed using the proposed multi-stack velocity-encoding scheme Velocity profiles taken across two tubes at the locations shown in (c) (inner diameters: P1 = 4mm and P2 = 8mm). Dashed lines correspond to PC-SPGR and solid lines correspond to bSSFP.

The 4D velocity vector reconstruction framework produced coherent flow patterns in all fetal cases. For evaluation of this data, the 4D magnitude cine volumes were used to orientate the heart in any arbitrary view enabling both through-plane and in-plane examination of blood flow through specific vessels or regions of the heart. There is a large amount of data in each subject, so one exemplar normal subject (ID03, GA24^+4^) will be described in detail. Figure 6 shows three different oblique cardiac views during systole and velocity-component volumes from the 4D flow cine reconstruction. In Figure 6a, the directionality of measured blood flow changes from positive to negative as the blood passes up the ascending aorta and through the descending aorta. A horizontal component of blood flow can be seen at the transverse arch of the aorta. In Figure 6b, increased blood flow can be seen where the superior vena cava joins the right atrium. In the third velocity-component volume (V^3^) of Figure 6c, blood in the aorta and pulmonary artery can be seen flowing with the same directionality.

**Figure 6:**
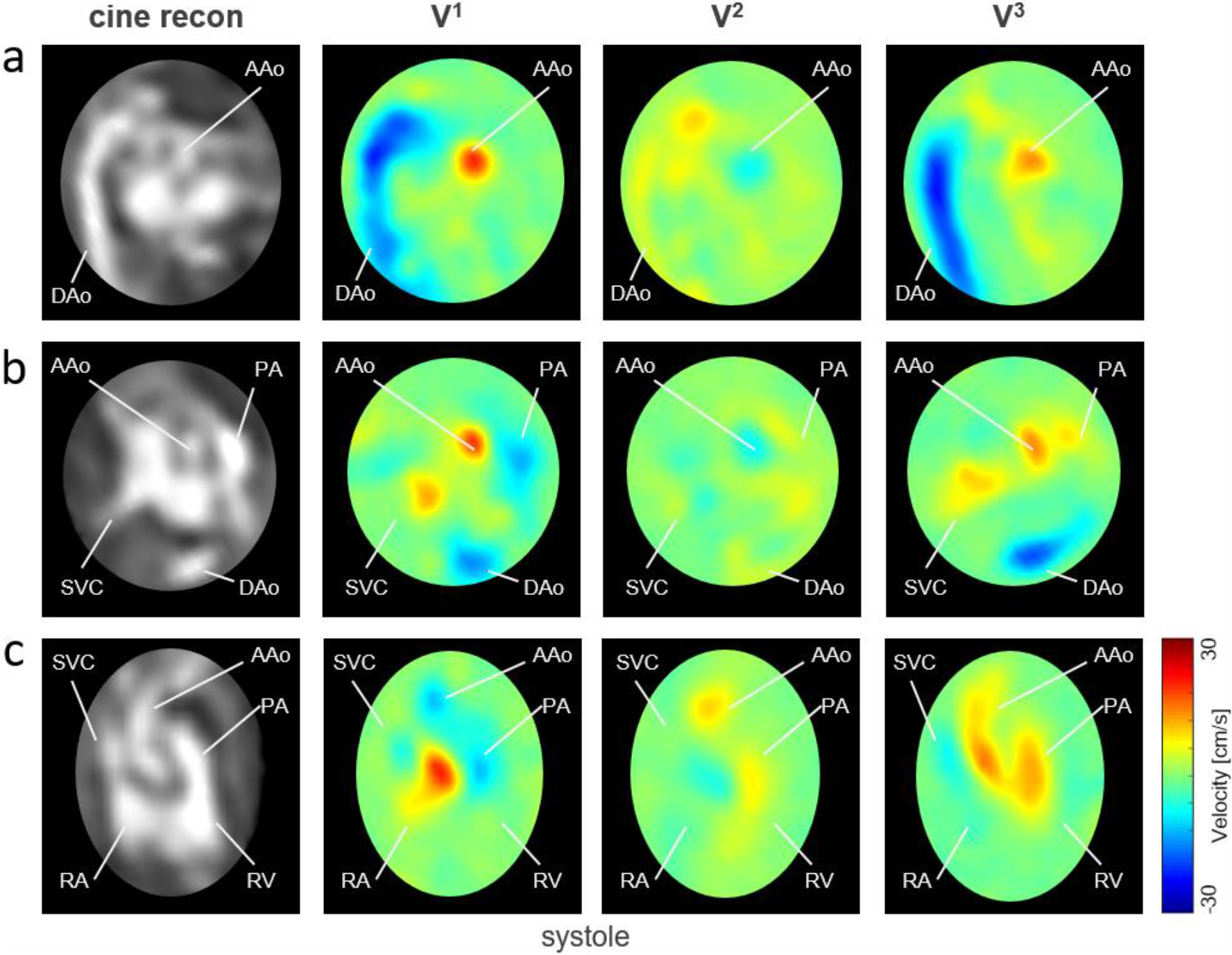
Three different slices at systole taken from 4D magnitude volume and corresponding velocity-component maps in subject ID03 (healthy fetus, 24^+2^ weeks GA). Magnitude images (left column) are shown with corresponding maps showing the three orthogonal components of the velocity field (V^1^, V^2^, V^3^). Rows: (a) Aortic arch plane: increased and opposing velocity can be seen in the ascending aorta (AAo) and descending aorta (DAo). Anti-parallel velocity can be seen between the AAo and DAo in components V^1^ and V^3^. (b) View showing the AAo, DAo, pulmonary artery (PA) and superior vena cava (SVC). Increased velocity is seen in all vessels compared to background velocity. (c) Three vessel view including the right chambers showing the AAo, PA, SVC, right atrium (RA) and right ventricle (RV). Velocities with common directionality can be seen in component V^3^, while reduced, antiparallel velocities are seen in the SVC.

4D flow cine volumes were generated by vector combination of the reconstructed velocity component volumes. Both 2D (Figure 7) and 3D visualisation (Figure 8) of velocity vectors reveal patterns of blood flow consistent with fetal hemodynamics. Pulsatile blood flow through the full extent of the aorta could be visualised through the course of the cardiac cycle (Supplementary Video S1). During systole, blood flow is fastest in the ascending aorta and descending aorta (Figure 7a and Figure 8a). Long-axis views the heart (Figure 7b and Figure 8b, Supplementary Video S2) show simultaneous outflow from the left- and right-ventricles through the left- and right outflow tracts. 3D visualisation demonstrates the normal spiral relationship of the outflow tracts (Figure 8b) and the full path of blood flow from the left ventricle through the left outflow tract and down the descending aorta. Lower magnitude patterns of blood flow can be visualised throughout the heart. Figure 7c and Figure 8c show right atrial inflow from the superior vena cava and inferior vena cava. In Figure 8c and Supplementary Video S3, this blood flow can be seen exiting the ventricle via the pulmonary artery. Cross-atrial blood flow through the foramen ovale can also be seen in Figure 7d.

**Figure 7:**
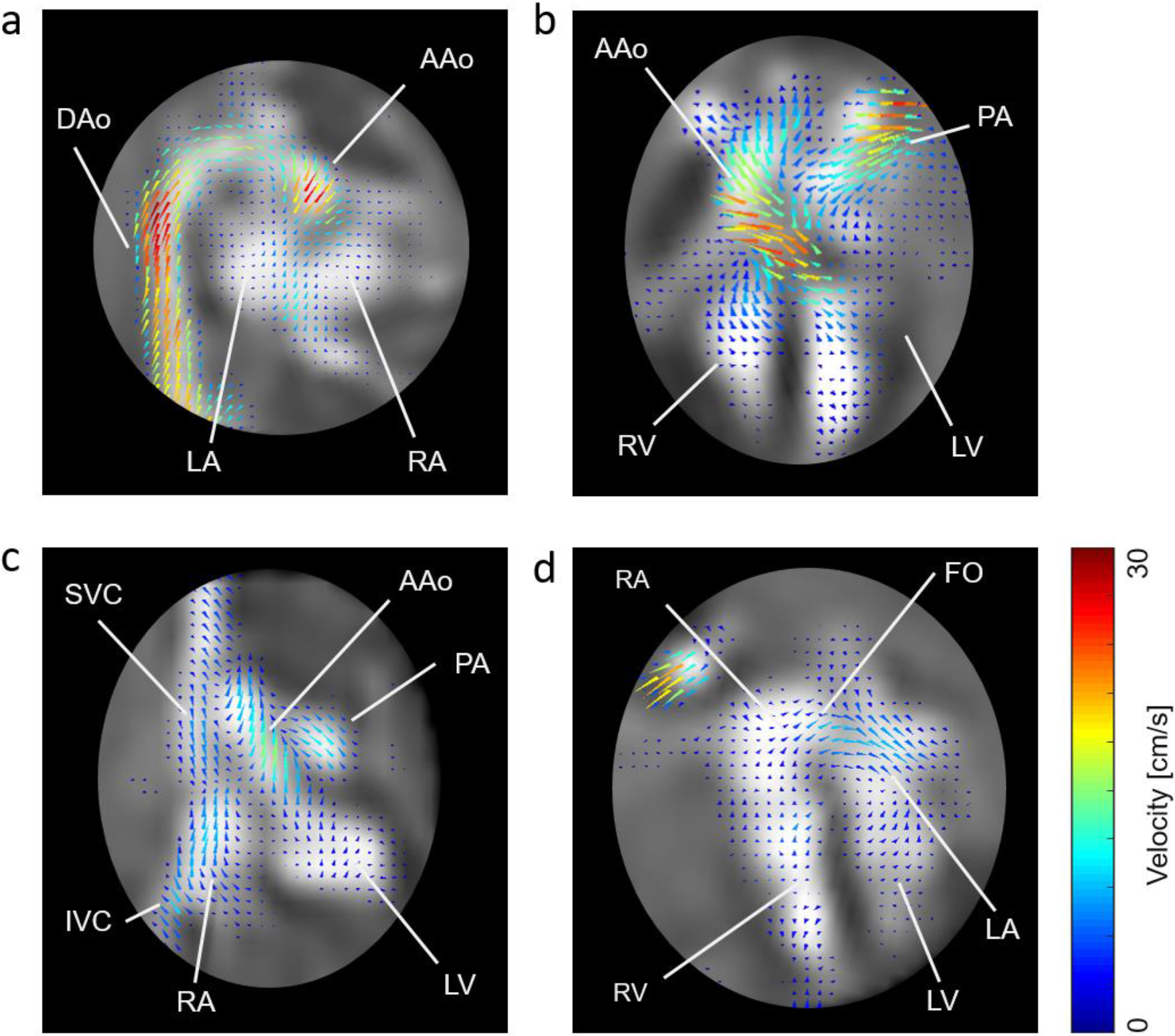
Velocity vector plots in subject ID03 (healthy fetus 24^+2^ weeks gestational age) overlaid on corresponding 4D magnitude cine frames. (a) Aortic arch plane showing blood flow vectors curve around the transverse arch of the aorta and down the descending aorta (DAo). (b) Dual-ventricle view showing blood flow vectors emerging from the left ventricle (LV) and right ventricle (RV) and passing through the ascending aorta (AAo) and pulmonary artery (PA), respectively. (c) Four-vessel view showing vectors flowing from the superior vena cava (SVC) and inferior vena cava (IVC) entering the right atrium (RA), and vectors passing through the AAo and PA. (d) Four-chamber view showing vector blood flow pass between the atria across the foramen ovale (FO).

**Figure 8:**
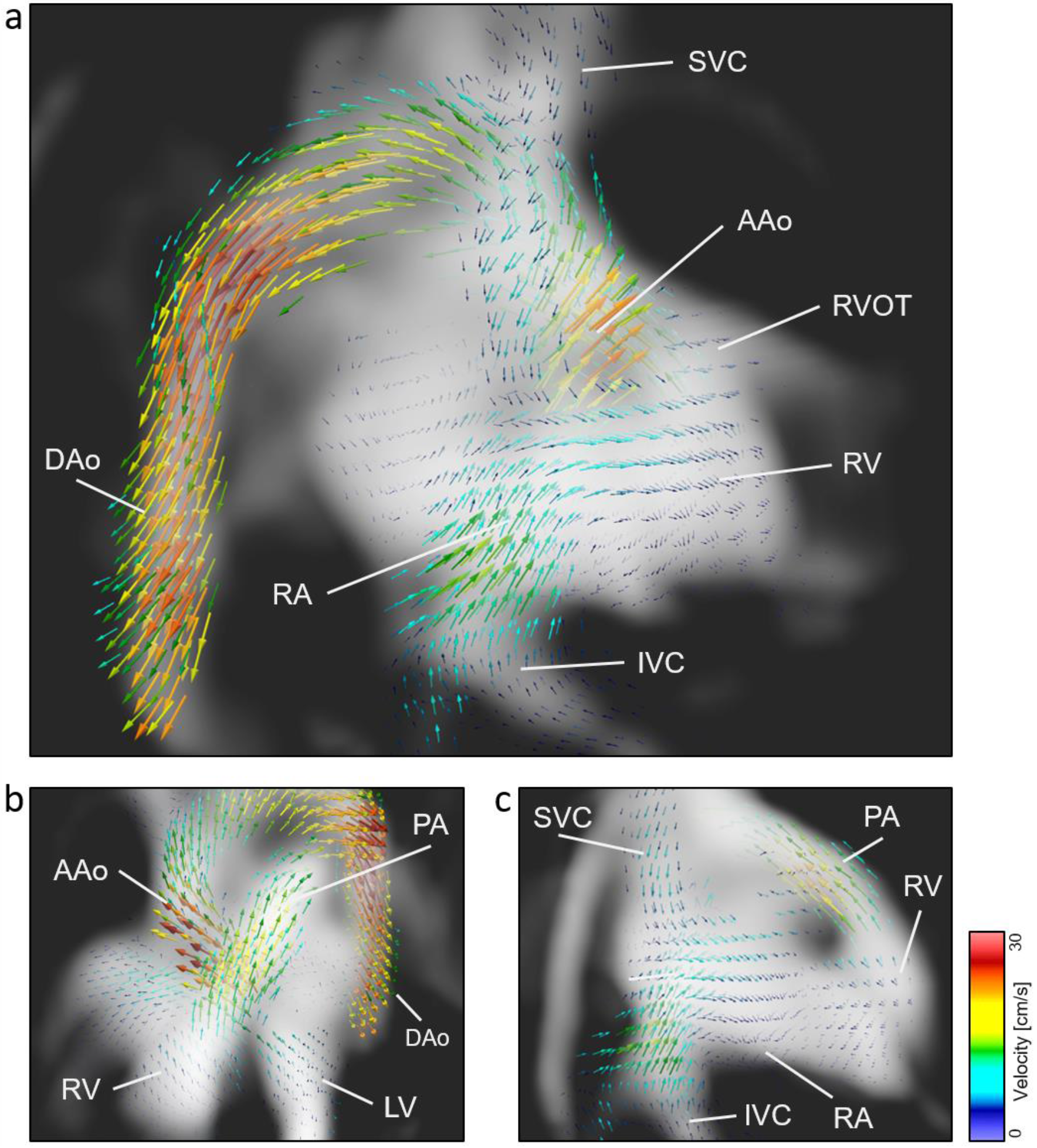
Volumetric renders showing 4D flow velocity vectors in subject ID03 (healthy fetus 24^+2^ weeks gestational age) overlaid on 4D magnitude cine renders, during atrial diastole. (a) Aortic arch view showing vectors emerging from the ascending aorta (AAo), flowing around the arch and through the descending aorta (DAo). (b) Long-axis view showing blood flow through the inter-crossing AAo and pulmonary artery (PA). (c) Right-sided view showing right atrium (RA) inflow from the superior vena cava (SVC) and inferior vena cava (IVC), which subsequently mixes and passes through to the right ventricle (RV) before flowing out of the pulmonary artery (PA). See supplementary material for videos of the views in this figure.

Plotting velocity data from the five measured vessels demonstrated expected pulsatile flow in the major arteries (the aorta, pulmonary artery, arterial duct and descending aorta) in the majority of cases. In all cases, the superior vena cava had a more constant flow curve throughout the cardiac cycle, in keeping with venous return flow. Figure 9a shows blood flow curves from fetus ID03 (healthy, GA 24^+4^). When taking the mean of the entire cohort (n = 7, Figure 9b), blood flow values were lower than expected from literature, by approximately a factor of 2 depending on the vessel. The pulmonary artery had the largest flow rate (138 ± 33 ml/min/kg), followed by the descending aorta (119 ± 56 ml/min/kg), ascending aorta (108 ± 31 ml/min/kg), ductus arteriosus (77 ± 54 ml/min/kg) and superior vena cava (74 ± 20 ml/min/kg).

**Figure 9:**
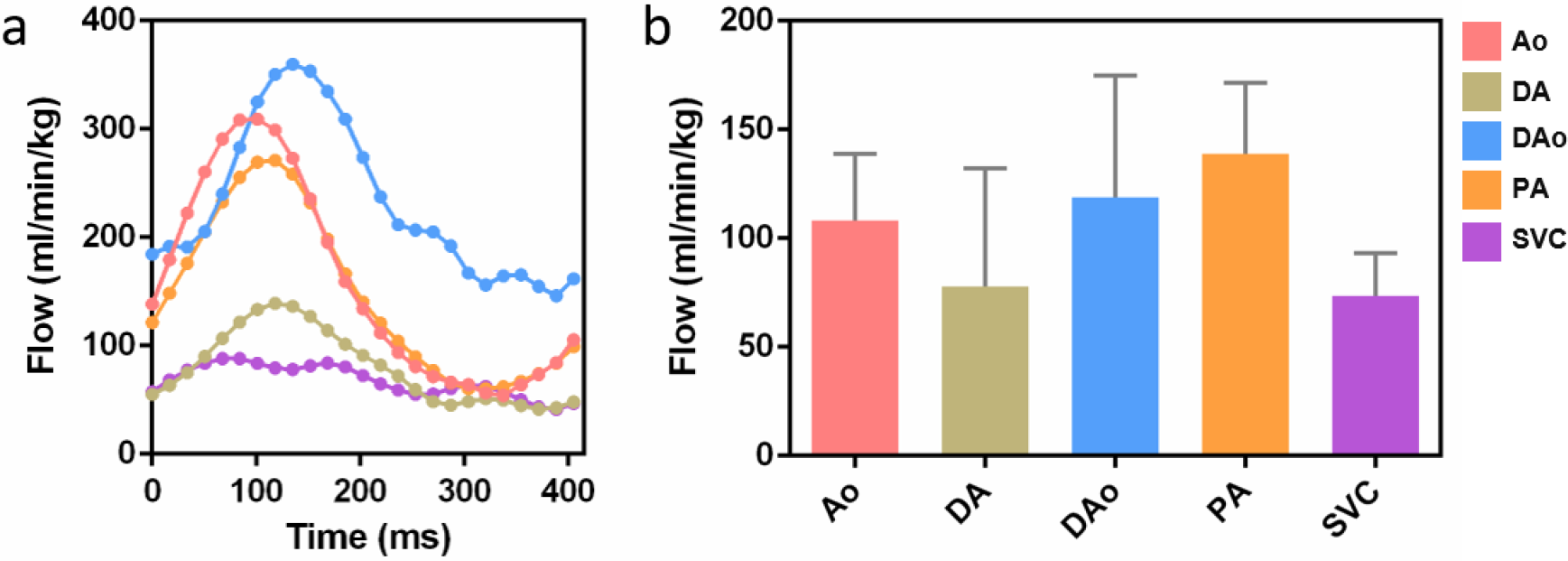
Analysis of fetal blood flow through the major vessels. (a) Flow curves measured in subject ID03 (healthy fetus, 24^+2^ weeks GA). The flow curves were derived using 2D cross-sectional regions of interest drawn by an expert fetal cardiologist on the 4D magnitude volume reconstructions, which were applied to the 4D flow reconstructed volumes. (b) Mean blood flows (+SD) across all seven fetal subjects. Values are lower than previously published literature values by a factor of approximately 2. Ao = aorta, DA = ductus arteriosus, DAo = descending aorta, PA = main pulmonary artery, SVC = superior vena cava.

## 5 DISCUSSION

In this work, we present a method for *in utero* human whole-heart fetal 4D cine quantitative blood flow imaging. 4D vector flow cine MRI reconstruction was achieved by extending our previous volumetric 4D magnitude cine framework^23^ to exploit the velocity-sensitive information inherent to the phase of dynamic bSSFP acquisitions. Multiple, non-coplanar bSSFP stacks were used to reconstruct spatially-identical, temporally-resolved, motion-corrected magnitude and vector flow volumes. Blood flow through fetal vasculature, the cardiac chambers and any other regions of interest could be inspected by re-orientation of 3D renders into any view or re-slicing volumes into any desired 2D projection. The method was validated using a digital phantom and directly compared with conventional PC-SPGR quantitative flow measurements using a simple constant flow phantom consisting of coiled tubing. Directionality of flows and absolute values measured using the proposed bSSFP method were found to be correct in both the digital and physical phantoms. Blood flow results obtained in the fetal subjects underestimated reference measurements in late-gestational fetuses^33,34^ by a factor of approximately two depending on the vessel of interest, however, the relative absolute flow rates of the vessels were consistent. There are numerous potential contributing factors including limited spatial and temporal resolution, as well as possible errors in background phase and velocity corrections.

One of the primary challenges of this work was reconstructing velocity measurements using phase contrast MRI without maintaining a fixed imaging volume between different velocity-encoding acquisitions. Volumetric reconstruction using slice-to-volume registration techniques^17–21^ requires acquisition of non-coplanar stacks of images, therefore a novel approach to velocity-encoding was developed. Previous work demonstrated that quantitative flow imaging was feasible using phase images from bSSFP sequences^26^. Nielsen et al.^27^ showed that multi-dimensional velocity-encoding could be achieved using balanced sequences by manipulating the gradient waveforms to impart orthogonal velocity-encoding, a concept which served as inspiration to combine multiple unique velocity-encoding samples associated with different multi-planar stacks. However, rather than manipulating the gradient waveforms to sample velocity-space, instead, identical gradient waveforms were used for all the acquired image stacks and 3D velocity-encoding was achieved purely through appropriate re-orientation of the imaging volumes and the velocity sensitive gradient moments associated with these.

Here, all experiments were performed using a minimum of five image stacks. In theory, 3D velocity encoding could be achieved using three stacks, however, fetal motion could then lead to local gaps in the sampled data that would compromise the inversion from component phases to full velocity, leading to inaccurate estimation of the final velocity vectors. Therefore, for this first demonstration of the proposed method we always required at least 5 image stacks with different orientations to ensure sufficient redundancy in the event of such motion. So far no attempt has been made to assess what is practically required of the imaging stacks, or what mitigation is appropriate to avoid incorrect results.

The physical flow phantom, which consisted of two tubes of flowing water with different internal diameters, demonstrated that the principles of non-coplanar velocity-encoding worked in practice and that flow measurements using a bSSFP sequence were accurate and directionally aligned with a gold-standard PC-SPGR acquisition. Measurement of peak velocity magnitude in the two tubes using PC-bSSFP slightly underestimated the PC-SPGR values, by a small difference of 8%, in both tubes.

For this exploratory work, a total of seven fetal cases were reconstructed including three healthy subjects, two right-sided aortic arch (RAA) cases and two further abnormal subjects, ranging from 24- to 32-weeks gestational age. 4D flow cine volumes were successfully reconstructed in all subjects, with pulsatile blood flow measured in all cases, consistent velocity magnitudes (ranging between 0 and an upper limit of 70 cm/s) between reconstructions, fast blood flow through the aorta and pulmonary artery and slower blood flow in the inferior vena cava, superior vena cava and the chambers of the hearts.

The background correction of the bSSFP phase images, demonstrated by Nielsen et al.^27^ and implemented here, was imperative for accurate inversion and calculation of velocity vectors. The uterus is particularly amendable to this method of phase correction because amniotic fluid fills the fetal lungs and surrounds the fetal body. This helps to minimise bSSFP-related artefacts, which would be caused particularly by air in the lungs of adults, as well as neonatal and pediatric subjects. Imaging at a relatively low 1.5T-field strength helps to reduce phase variation as well. The proposed framework is field-strength independent, but the bSSFP acquisition will need optimisation and testing at different field strengths.

Quantification of blood flow curves in the major vessels was carried out using simple 2D cross-sectional regions of interest, in keeping with current PC-MRI blood flow methods^13,14^, however, 3D segmentation of the fetal heart could improve flow quantification. In comparison with single-slice metric-optimised gating (MOG) fetal blood flow techniques^33,34^, the absolute blood flow values measured here were lower in all vessels by a factor of approximately two. The relative absolute flow rates were in in keeping with literature values: flow was fastest in the pulmonary artery, followed by the descending aorta, ascending aorta, ductus arteriosus and the superior vena cava. Comparatively, blood flow curves in the ductus arteriosus were still low despite the relative consistency. The reasons for this are not immediately clear, although the arterial duct was consistently reported as the most difficult vessel to identify within the resolution limits of 3D magnitude images. This is the subject of ongoing investigation.

The overall directionality of the 4D vector flow cine volumes and temporal fluctuations through the cardiac cycle provide confidence that the underlying principles of the framework are correct. However, some degree of intra-subject variability was observed in some small localised regions, where velocity vectors pointed in unexpected orientations, such as perpendicular to the anticipated direction of flow. A range of issues could explain this including incomplete velocity-encoding due to fetal and cardiac motion, or regions not sampled by a sufficient fraction of the phase stacks. The low absolute values of the blood flow curves can be attributed to the constraints of the bSSFP acquisition, which are relatively low spatial resolution (2.0×2.0×6.0 mm) and relatively long temporal resolution (72 ms), compared to methods using MOG^33^ (1.25-1.25×5.0 mm and 50 ms). An acquisition scheme tailored to 4D flow cine imaging will be the focus of future work, to improve spatial and temporal resolution. Despite these limitations, the acquisition protocol brings significant advantages compared to single-slice methods as it eliminates the need for high precision slice plane selection during planning which is challenging and prone to error. The proposed method also allows for simultaneous structural imaging and complete flow mapping, enabling contemporaneous comparisons of flow patterns at diverse locations in the fetal circulatory system, which are more clinically meaningful. The potential for studying flow could be extended to intra-cardiac flow patterns, which could enhance understanding of perturbations in the setting of structural lesions.

## CONCLUSION

A fully motion-corrected 4D vector flow cine imaging method for fetal cardiac visualisation based on accelerated bSSFP sequences has been presented. The method was validated using simulations and using a simple constant flow phantom before being deployed in seven fetal subjects. Results confirm that the method is robust, provides coherent temporally- and spatially-resolved vector velocity fields, and can yield quantitative flow data for major fetal vessels. Despite validation data that confirms accuracy for steady flows in phantoms, at present measured flow rates in fetal vessels are lower than would be expected. There are many opportunities for improving and refining the method and these will be the subject of future research.

## ACKNOWLEDGEMENT

Thank you to Joanna Allsop, Elaine Green and Ana Gomes for scanning of volunteers and patients.

## REFERENCES

1. Kenny JF, Plappert T, Doubilet P, Saltzman DH, Cartier M, Zollars L, Leatherman G, St John Sutton M. Changes in intracardiac blood flow velocities and right and left ventricular stroke volumes with gestational age in the normal human fetus: a prospective Doppler echocardiographic study. Circulation 1986;74(6):1208–1216.

2. Hecher K, Campbell S, Doyle P, Harrington K, Nicolaides K. Assessment of fetal compromise by Doppler ultrasound investigation of the fetal circulation: arterial, intracardiac, and venous blood flow velocity studies. Circulation 1995;91(1):129–138.

3. Arabin B. Doppler blood flow measurement in uteroplacental and fetal vessels: pathophysiological and clinical significance: Springer Science & Business Media; 2012.

4. Blanco P. Volumetric blood flow measurement using Doppler ultrasound: concerns about the technique. Journal of ultrasound 2015;18(2):201–204.

5. Markl M, Frydrychowicz A, Kozerke S, Hope M, Wieben O. 4D flow MRI. Journal of Magnetic Resonance Imaging 2012;36(5):1015–1036.

6. Lotz J, Meier C, Leppert A, Galanski M. Cardiovascular flow measurement with phase-contrast MR imaging: basic facts and implementation. Radiographics 2002;22(3):651–671.

7. Nayak KS, Nielsen J-F, Bernstein MA, Markl M, Gatehouse PD, Botnar RM, Saloner D, Lorenz C, Wen H, Hu BS. Cardiovascular magnetic resonance phase contrast imaging. Journal of Cardiovascular Magnetic Resonance 2015;17(1):71.

8. Lawley CM, Broadhouse KM, Callaghan FM, Winlaw DS, Figtree GA, Grieve SM. 4D flow magnetic resonance imaging: role in pediatric congenital heart disease. Asian Cardiovascular and Thoracic Annals 2018;26(1):28–37.

9. Powell A, Maier S, Chung T, Geva T. Phase-velocity cine magnetic resonance imaging measurement of pulsatile blood flow in children and young adults: in vitro and in vivo validation. Pediatric cardiology 2000;21(2):104–110.

10. Hsiao A, Alley MT, Massaband P, Herfkens RJ, Chan FP, Vasanawala SS. Improved cardiovascular flow quantification with time-resolved volumetric phase-contrast MRI. Pediatric radiology 2011;41(6):711–720.

11. Broadhouse KM, Price AN, Durighel G, Cox DJ, Finnemore AE, Edwards AD, Hajnal JV, Groves AM. Assessment of PDA shunt and systemic blood flow in newborns using cardiac MRI. Nmr Biomed 2013;26(9):1135–1141.

12. Broadhouse KM, Price AN, Finnemore AE, Cox DJ, Edwards AD, Hajnal JV, Groves AM. 4D phase contrast MRI in the preterm infant: visualisation of patent ductus arteriosus. Archives of Disease in Childhood-Fetal and Neonatal Edition 2015;100(2):F164–F164.

13. Jansz MS, Seed M, van Amerom JF, Wong D, Grosse-Wortmann L, Yoo SJ, Macgowan CK. Metric optimized gating for fetal cardiac MRI. Magnetic resonance in medicine 2010;64(5):1304–1314.

14. Goolaub DS, Roy CW, Schrauben E, Sussman D, Marini D, Seed M, Macgowan CK. Multidimensional fetal flow imaging with cardiovascular magnetic resonance: a feasibility study. Journal of Cardiovascular Magnetic Resonance 2018;20(1):77.

15. Kording F, Schoennagel BP, Ruprecht C, Giese D, Heiberg E, Tavares M, Yamamura J. Feasibility of 4D phase-contrast MRI for the assessment of blood flow in the fetal aorta using Doppler ultrasound gating: preliminary results. 2018.

16. Schrauben EM, Saini BS, Darby JR, Soo JY, Lock MC, Stirrat E, Stortz G, Sled JG, Morrison JL, Seed M. Fetal hemodynamics and cardiac streaming assessed by 4D flow cardiovascular magnetic resonance in fetal sheep. Journal of Cardiovascular Magnetic Resonance 2019;21(1):8.

17. Kuklisova-Murgasova M, Quaghebeur G, Rutherford MA, Hajnal JV, Schnabel JA. Reconstruction of fetal brain MRI with intensity matching and complete outlier removal. Medical image analysis 2012;16(8):1550–1564.

18. Gholipour A, Estroff JA, Warfield SK. Robust super-resolution volume reconstruction from slice acquisitions: application to fetal brain MRI. IEEE transactions on medical imaging 2010;29(10):1739–1758.

19. Jiang S, Xue H, Glover A, Rutherford M, Rueckert D, Hajnal JV. MRI of moving subjects using multislice snapshot images with volume reconstruction (SVR): application to fetal, neonatal, and adult brain studies. IEEE transactions on medical imaging 2007;26(7):967–980.

20. Rousseau F, Kim K, Studholme C, Koob M, Dietemann J-L. On super-resolution for fetal brain MRI. 2010. Springer. p 355–362.

21. Lloyd DF, Pushparajah K, Simpson JM, van Amerom JF, van Poppel MP, Schulz A, Kainz B, Deprez M, Lohezic M, Allsop J. Three-dimensional visualisation of the fetal heart using prenatal MRI with motion-corrected slice-volume registration: a prospective, single-centre cohort study. The Lancet 2019;393:1619–1627.

22. van Amerom JF, Lloyd DF, Price AN, Kuklisova Murgasova M, Aljabar P, Malik SJ, Lohezic M, Rutherford MA, Pushparajah K, Razavi R. Fetal cardiac cine imaging using highly accelerated dynamic MRI with retrospective motion correction and outlier rejection. Magnetic resonance in medicine 2018;79(1):327–338.

23. van Amerom JF, Lloyd DF, Deprez M, Price AN, Malik SJ, Pushparajah K, van Poppel MP, Rutherford MA, Razavi R, Hajnal JV. Fetal whole-heart 4D imaging using motion-corrected multi-planar real-time MRI. Magnetic resonance in medicine 2019. doi: 10.1002/mrm.27798.

24. Tsao J, Boesiger P, Pruessmann KP. k-t BLAST and k-t SENSE: dynamic MRI with high frame rate exploiting spatiotemporal correlations. Magnetic Resonance in Medicine: An Official Journal of the International Society for Magnetic Resonance in Medicine 2003;50(5):1031–1042.

25. Tsao J, Kozerke S, Boesiger P, Pruessmann KP. Optimizing spatiotemporal sampling for k-t BLAST and k-t SENSE: application to high-resolution real-time cardiac steady-state free precession. Magnetic Resonance in Medicine: An Official Journal of the International Society for Magnetic Resonance in Medicine 2005;53(6):1372–1382.

26. Markl M, Alley M, Pelc N. Balanced phase-contrast steady-state free precession (PC-SSFP): a novel technique for velocity encoding by gradient inversion. Magnetic Resonance in Medicine: An Official Journal of the International Society for Magnetic Resonance in Medicine 2003;49(5):945–952.

27. Nielsen JF, Nayak KS. Referenceless phase velocity mapping using balanced SSFP. Magnetic Resonance in Medicine: An Official Journal of the International Society for Magnetic Resonance in Medicine 2009;61(5):1096–1102.

28. Deprez M, Price AN, Christiaens D, Estrin GL, Grande LC, Hutter J, Daducci A, Tournier J-D, Rutherford MA, Counsell S. Higher order spherical harmonics reconstruction of fetal diffusion MRI with intensity correction. bioRxiv 2018:297341.

29. Hand J, Li Y, Hajnal J. Numerical study of RF exposure and the resulting temperature rise in the foetus during a magnetic resonance procedure. Physics in Medicine & Biology 2010;55(4):913.

30. Glover P, Hykin J, Gowland P, Wright J, Johnson I, Mansfield P. An assessment of the intrauterine sound intensity level during obstetric echo-planar magnetic resonance imaging. The British journal of radiology 1995;68(814):1090–1094.

31. Tournier J-D, Smith RE, Raffelt DA, Tabbara R, Dhollander T, Pietsch M, Christiaens D, Jeurissen B, Yeh C-H, Connelly A. MRtrix3: A fast, flexible and open software framework for medical image processing and visualisation. bioRxiv 2019:551739.

32. Hadlock FP, Harrist RB, Martinez-Poyer J. In utero analysis of fetal growth: a sonographic weight standard. Radiology 1991;181(1):129–133.

33. Prsa M, Sun L, van Amerom J, Yoo S-J, Grosse-Wortmann L, Jaeggi E, Macgowan C, Seed M. Reference ranges of blood flow in the major vessels of the normal human fetal circulation at term by phase-contrast magnetic resonance imaging. Circulation: Cardiovascular Imaging 2014;7(4):663–670.

34. Seed M, van Amerom JF, Yoo S-J, Al Nafisi B, Grosse-Wortmann L, Jaeggi E, Jansz MS, Macgowan CK. Feasibility of quantification of the distribution of blood flow in the normal human fetal circulation using CMR: a cross-sectional study. Journal of cardiovascular magnetic resonance 2012;14(1):79.

