## Supplementary figures and images for "Fetal whole-heart 4D flow cine MRI using multiple non-coplanar balanced SSFP stacks"

### Supplementary Figure 1

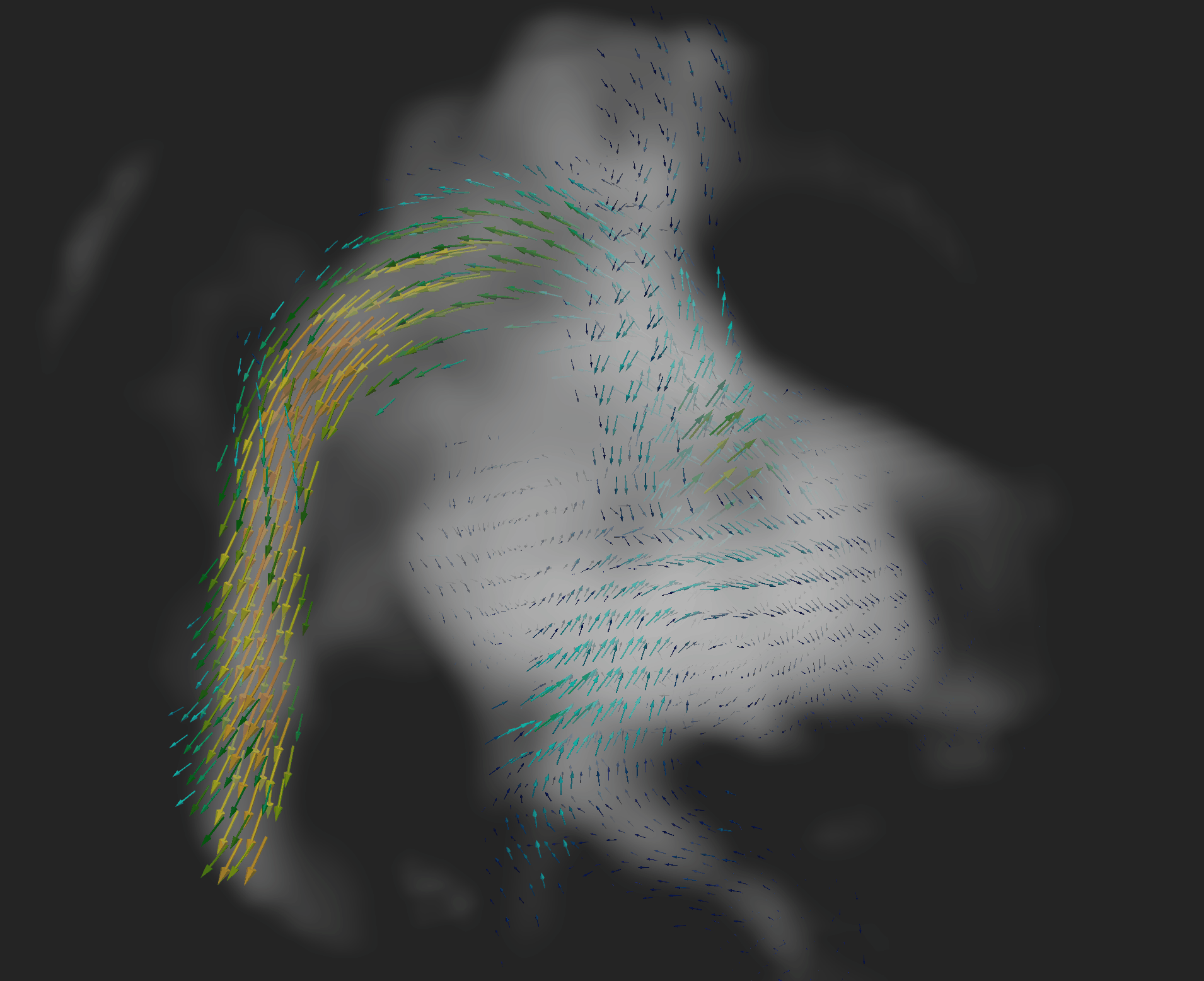

### Supplementary Figure 2

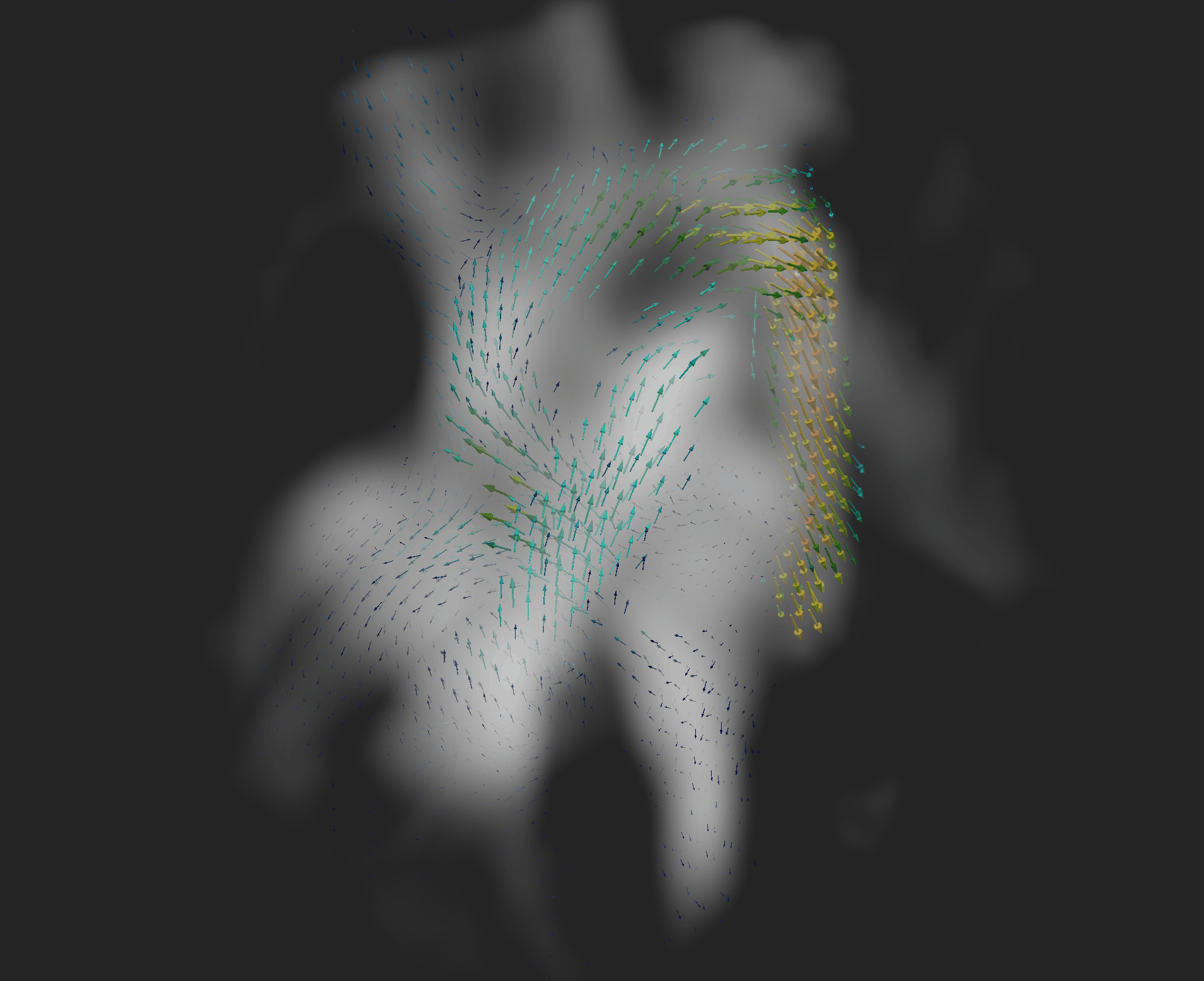

### Supplementary Figure 3

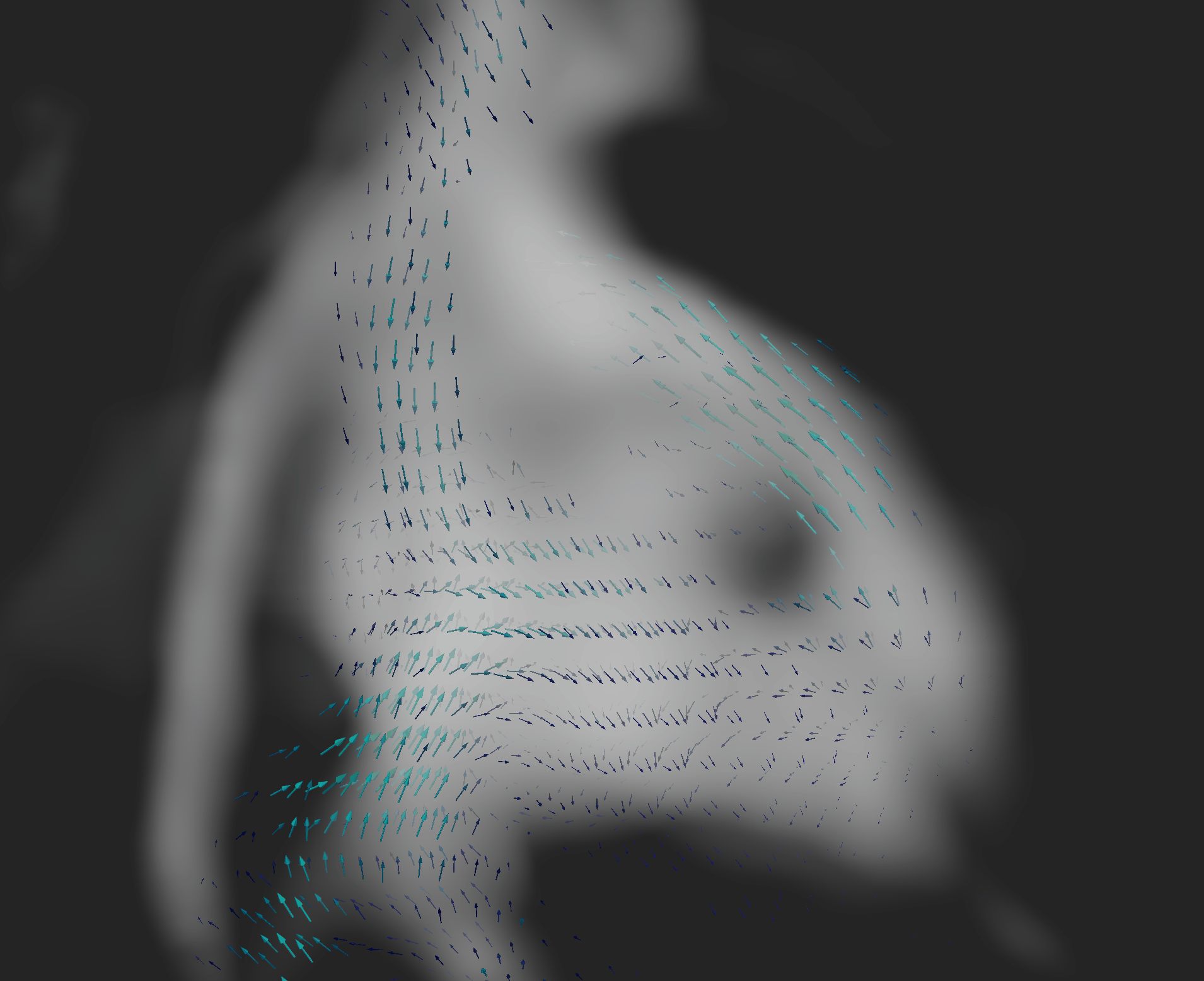
